# Comprehensive palmitoyl-proteomic analysis identifies distinct protein signatures for large and small cancer-derived extracellular vesicles

**DOI:** 10.1101/787499

**Authors:** Javier Mariscal, Tatyana Vagner, Minhyung Kim, Bo Zhou, Andrew Chin, Mandana Zandian, Michael R. Freeman, Sungyong You, Andries Zijlstra, Wei Yang, Dolores Di Vizio

**Author notes:** Corresponding authors: Dolores Di Vizio, Wei Yang, Andries Zijsltra.

## Abstract

Extracellular vesicles (EVs) are membrane-enclosed particles that play an important role in cancer progression and have emerged as a promising source of circulating biomarkers. Protein *S*-acylation, also known as palmitoylation, has been proposed as a post-translational mechanism that modulates the dynamics of EV biogenesis and protein cargo sorting. However, technical challenges have limited large-scale profiling of the whole palmitoyl-proteins of EVs. We successfully employed a novel approach that combines low-background acyl-biotinyl exchange (LB-ABE) with label-free proteomics to analyze the palmitoyl proteome of large EVs (L-EVs) and small EVs (S-EVs) from prostate cancer cells. Here we report the first palmitoyl-protein signature of EVs, and demonstrate that L- and S-EVs harbor proteins associated with distinct biological processes and subcellular origin. We identified STEAP1, STEAP2, and ABBC4 as prostate cancer-specific palmitoyl proteins enriched in both EV populations in comparison with the originating cell lines. Importantly, the presence of the above proteins in EVs was significantly reduced upon inhibition of palmitoylation in the producing cells. These results suggest that palmitoylation may be involved in the differential sorting of proteins to distinct EV populations and allow for better detection of disease biomarkers.

## Introduction

Protein *S*-acylation, more commonly known as *S*-palmitoylation, and hereafter referred to as palmitoylation, is a reversible post-translational modification (PTM) where long-chain fatty acids are covalently attached to cysteine residues via labile thioester bonds^1^. Palmitoylation increases protein hydrophobicity, transiently targeting cytosolic proteins to cell membranes. Among all lipid modifications, palmitoylation is the most pervasive, affecting about 20% of the proteome^2,3^. Additionally, palmitoylation tethers proteins into membrane microdomains such as lipid rafts, thus modulating protein activity, stability, and multiprotein complex formation^4^. Palmitoylation is frequently altered in various diseases including cancer^3,5^. However, the biological role of this PTM is complex and multifaceted and has not been fully investigated due to technical challenges associated with the low abundance of palmitoyl-proteins in general, the difficulty of enriching palmitoyl-proteins with high specificity, and the high hydrophobicity of intact palmitoyl-peptides.

Extracellular vesicles (EVs) are lipid-enclosed particles that play an important role in cancer progression and have emerged as a promising source of circulating biomarkers. Several classes of EVs that differ in size, cargo, biogenesis and function have been reported in the last decade^6–10^. However, clearly defined markers that define these classes across different cell lines are currently lacking^11^. To address this challenge the International Society for Extracellular Vesicles has proposed that EV should be classified into two major populations of vesicles based on their size: small EVs (S-EVs) and large EVs (L-EVs). S-EVs usually include vesicles in the 40–150 nm size range while L-EVs include vesicles in the 200 nm–10 μm size range^11^. However, these size ranges are not precise and can vary depending on the model system and measurement approach. S-EVs are represented both by exosomes, which originate from endocytic machinery and by small ectosomes, which originate from membrane blebs pinching off the plasma membrane^12,13^. However, endosomal origin of S-EVs is difficult to prove and most common isolation methods do not separate endosome-derived S-EVs from non-endosomal S-EVs^14–18^. In contrast, L-EVs frequently form at the plasma membrane by direct shedding of membrane blebs from the cells, and are represented by microvesicles, large oncosomes, apoptotic bodies, and other types of vesicles, mostly in the micrometer size range^19^. Several pathways involved in regulating the biogenesis of these two EV classes have been proposed^12^, but a lot is still unknown.

Several laboratories have used mass spectrometry to profile EVs^20,21^. These studies have identified numerous proteins with differential expression levels, not only between EVs and their cells of origin but also between different EV populations. We and others have shown that L- and S-EVs originating from the same cell share a large number of proteins, however there are also proteins that are differentially expressed between L-EVs and S-EVs^8^. For instance, integrin and adhesion molecules are enriched in S-EVs while metabolic enzymes and structural cytoplasmic proteins are enriched in L-EVs^8^. Despite these findings, EV proteomics has proven to be challenging due to co-isolation of non-EV proteins such as lipoproteins during EV isolation, which can mask the presence of less abundant, but potentially physiologically important, EV proteins. Palmitoylation can target proteins to the membrane of EVs^22^, suggesting that palmitoylated proteins are enriched in EVs. However, a recent study that analyzed PTMs of exosomes in depth^23^ was conducted using a conventional proteomics procedure, and in that case palmitoylation was not detectable because the thioester bonds were cleaved off during the sample preparation and the palmitoyl groups were lost. Consequently, palmitoylated cysteines could not be distinguished from non-palmitoylated cysteines.

A whole palmitoyl-proteomics analysis of EVs has not yet been performed. We employed a novel method that uses a metabolic labeling-independent, cysteine centric approach, namely low background acyl-biotin exchange (LB-ABE) to the study of EV palmitoyl-proteome. We sought to determine if label-free proteomics combined with LB-ABE can identify specific palmitoyl-proteomic signatures in EVs by comparing the palmitoyl-proteome of L- and S-EVs versus their parental cells. We also investigated whether these signatures reflect a specific subcellular origin and/or biological function. We also attempted to identify prostate cancer specific palmitoyl-proteins that are highly abundant in EVs.

## Methods

### Cell culture

The PC3 cell line was obtained from the American Type Culture Collection (ATCC). The DU145^DIAPH3-KD^ cell line, stably transfected with DIAPH3 shRNA, was generated in our laboratories^24^. PC3 and DU145^DIAPH3-KD^ cell lines were cultured in DMEM (Invitrogen). All cells were supplemented with 10% fetal bovine serum (Denville Scientific), 2mM L-glutamine (Invitrogen) and 1% PenStrep (Invitrogen). DU145^DIAPH3-KD^ cells were additionally selected with 2 µg/mL puromycin as described^24^. All cells were grown at 37°C and 5% CO_2._ Cell viability of the EV-producer cells was tested with the 0.4% Trypan Blue solution (Sigma) dye exclusion method. All cell lines were routinely tested for mycoplasma contamination by using the MycoAlert PLUS Mycoplasma Detection Kit (Lonza). Finally, in order to collect EVs from cells in which palmitoylation was inhibited, PC3 cells were treated from 24 hours with 2-bromohexadecanoic acid (Sigma), also known as 2-bromopalmitate (0.5, 1, 5, 10, 20, 50 and -100 µM), in serum-starvation.

### Isolation of extracellular vesicles (EVs)

The isolation of EVs was conducted as previously described with minor modifications^7,8,25^. Cells were grown on 18 x 150 cm^2^-cell culture plates (Corning) until 90% confluence, washed in PBS, and serum-starved for 24 hours before the collection of conditioned cell media. The conditioned media was cleared by differential centrifugation of floating cells at 300*g*, of cell debris at 2,800*g* for 10 min, and spun in an ultracentrifuge at 10,000*g* for 30 min (4°C, k-factor 2547.2) for the collection of L-EVs. The supernatant was then spun at 100,000*g* for 60 min (4°C, k-factor 254.7) for the collection of S-EVs. Both L-EVs and S-EVs were then subjected to Optiprep(tm) (Sigma) density gradient purification. Fresh pelleted EVs were resuspended in 0.2 µm-filtered PBS and deposited at the bottom of an ultracentrifuge tube. Next, 30% (4.3 mL, 1.20 g/mL), 25% (3 mL, 1.15 g/mL), 15% (2.5 mL, 1.10 g/mL), and 5% (6 mL, 1.08 g/mL) iodixanol solutions were sequentially layered at decreasing density to form a discontinuous gradient. Separation was performed by ultracentrifugation at 100,000*g* for 3h 50 min (4°C, k-factor 254.7) and EV-enriched fractions collected either at 1.10-1.15 g/mL for L-EVs or 1.10 g/mL for S-EVs^8^. Purified EVs were then washed in PBS (100,000*g*, 60 min, 4°C) and resuspended in the appropriate buffer. All ultracentrifugation spins were performed in a SW28 swinging rotor (Beckman Coulter). We have submitted all relevant data of our experiments to the EV-TRACK knowledgebase (EV-TRACK ID: EV190069)^26^.

### Whole cell and membrane protein lysates from EV-producer cells

Whole cell lysate (WCL) and membrane preparations (M) were obtained upon 24-hour serum-starvation and collection of conditioned cell media. Cell monolayers were scraped and washed in chilled PBS (x3). For WCL, cells were directly lysed in DTT-free 4% SDS/Tris-HCl lysis buffer^27^. For M preparations, cells were gently scraped, washed in PBS (x3) and resuspended in filtered PBS containing 1% protease inhibitors (cOmplete Mini Protease Inhibitor Cocktail, Roche). Cell suspensions were immediately subjected to 20 cycles of sonication (5 sec) in ice to induce cell disruption. Membrane suspensions were then cleared of intact cells at 500*g*, pelleted at 16,000*g* (20 min, 4°C), washed in PBS (x3) and resuspended in 4% SDS/Tris-HCl lysis buffer^4^. All protein lysates were stored at -80°C until use.

### LB-ABE enrichment of palmitoyl-proteins

Protein concentration was determined with the Pierce 660nm protein assay (Pierce). 300 µg of WCL, M and L-EVs (x3); and 250 µg of S-EVs (x2) were subjected to LB-ABE coupled to label free mass spectrometry as described^2^. Briefly, proteins were reduced with 50 mM tris(2-carboxyethyl)phosphine (TCEP), sequentially alkylated with 50 mM N-ethylmaleimide (NEM) and 25 mM 2,2’-dithiodipyridine (DTDP), and biotinylated with 1 mM biotin-HPDP in the presence or absence of 2 M hydroxylamine (Hyd). Palmitoyl-proteins were enriched by streptavidin affinity purification, eluted by 50 mM TCEP, and precipitated by methanol/chloroform.

In order to evaluate the specificity of the LB-ABE method, 1 mg of WCL and M protein lysates were subjected to LB-ABE. 2 µg of the recovered palmitoylated fractions were resolved on a SDS-acrylamide gel alongside with 2 µg of the whole protein and the non-palmitoylated fractions. Finally, silver staining of the gel was performed with the Silver Stain Kit (Pierce) following manufacturer’s recommendations. Alternatively, for the validation of select palmitoyl-proteins in WCL, M, L-EVs and S-EVs, 300 µg of total protein per group were processed as above with minor modifications. Briefly, samples were split into two equal halves in order to constitute the experimental (Hyd+) and negative control (Hyd-) groups prior to LB-ABE chemistry^2^. Experimental groups were subjected to 2 M Hyd treatment as above, whereas control groups were incubated with Tris/HCl buffer in order to evaluate the unspecific recovery of non-palmitoylated proteins. Upon recovery of palmitoylated proteins, experimental and control samples were re-dissolved in loading buffer and 10% (v/v) of total recovered proteins was loaded onto SDS-PAGE gels for immunoblotting analysis.

### LC-MS/MS analysis and data processing

Enriched palmitoyl-proteins were digested with MS-grade trypsin (Promega) by filter-aided sample preparation (FASP) as described previously^2,28^. Tryptic peptides were then recovered, dried down in a SpeedVac concentrator (Thermo Scientific), and re-dissolved in 0.2% formic acid (Sigma) up to a concentration of 0.15 µg/mL. Label-free proteomic analysis was performed using an EASY-nLC 1000 connected to an LTQ Orbitrap Elite hybrid mass spectrometer essentially as we previously described^29,30^. Briefly, 7 μL of peptide solution was loaded onto a 2-cm trap column (75 μm × 2 cm, C_18_) and separated on a 50-cm EASY-Spray analytical column (PepMap RSLC C_18_, 2 μm, 100Å, 50 μm × 15 cm) heated to 55°C, using a 2-h gradient consisting of 2-40% B in 150 min, 40-100% B in 20 min, and 100% B in 10 min at the flow rate of 150 nL/min. Separated peptides were ionized with an EASY-Spray ion source. Mass spectra were acquired in a data-dependent manner, with automatic switching between MS and MS/MS scans. In MS scans, the lock mass at m/z 445.120025 was applied to provide real-time internal mass calibration. The full MS scan (400-1600 m/z) was performed in 240,000 resolution at m/z of 400 Th, with an ion packet setting of 1×10^6^ for automatic gain control and a maximum injection time of 500 ms. Up to 20 most intense peptide ions with charge state of ≥2 were automatically selected for MS/MS fragmentation by rapid collision-induced dissociation (rCID), using 7,500 resolution, 1×10^4^ automatic gain control, 50 ms maximum injection time, 10 ms activation time, and 35% normalized collision energy. To minimize redundant spectral acquisition, dynamic exclusion was enabled with a repeat count of 1, an exclusion during of 30 s, and a repeat duration of 60 s.

The acquired MS data were searched against the Uniprot_Human database (released on 01/22/2016, containing 20,985 protien sequences) using the Andromeda^31^ algorithm in theMaxQuant (v1.5.5.1)^32^ environment. The searching parameters were set as follows: trypsin/P as the protease; oxidation (M), acetyl (protein N-term), NEM(C), and carbamidomethyl(C) as variable modifications; up to two missed cleavages; minimal peptide length as 7; mass tolerance for MS1 was 4.5 ppm for main search and for MS2 was 0.5 Da; identification of second peptides enabled; label free quantification (LFQ) enabled, and match-between-runs within 2 min were enabled. A stringent <0.01 FDR was used to filter PSM, peptide, and protein identifications.

### Identification of high-confidence and high-abundance proteins

To retain high confidence proteins from the raw LFQ data, we applied criteria that proteins should be detected with at least 2 peptides and in at least 2 replicates and low abundant proteins less than 5% of absolute protein abundance for each identification were discarded for subsequent analyses. Absolute protein abundance for each identification was defined as the median value of replicates. We assumed that total protein amount and composition of protein species are different between WCL, M, L-EVs and S-EVs. To compare protein abundance between them, a max-min normalization method was used to scale the protein expression values between 0 to 1. Protein expression value 1 was given to the most abundant protein within each sample and protein value 0 was given to the least abundant protein identified. Finally, in order to detect significantly abundant proteins within each sample, the rank product algorithm was applied to the normalized protein abundance for each identification^33^. In order to identify high-abundance proteins in EVs, WCL or M, we performed integrated hypothesis test. Briefly, Student’s t-test and log_2_-median ratio test were performed. We estimated empirical null distributions of T values and log_2_-median ratio value by randomly permuting all samples 1,000 times and calculating t-test and log_2_-median ratio test p-values. We then integrated these two p-value into an overall P-value using Stouffer’s method^34^. False discovery rate (FDR) was corrected by Storey’s method^35^. We then selected proteins with FDR <0.05 and normalized expression difference ≥0.1. To set the normalized expression difference, we generated null hypothesis distribution by randomly permuting all samples 1,000 times and set the threshold at 99 percentile value of the distribution, which is 0.1.

### Functional annotation of the palmitoyl-proteome

The percentage of putative high-confidence (identified by 2 independent methods in palmitoyl-proteomes or experimentally validated) human palmitoyl-proteins in cells was directly retrieved from the SwissPalm database (v2) database (www.swisspalm.org)^36^. For EVs, the percentage of putative palmitoyl-proteins was estimated by direct comparison to the number of human proteins retrieved from the ExoCarta database (www.ExoCarta.org)^37^. To assess the biological relevance of the putative palmitoyl-proteins in the proteome of prostate cancer EVs, we selected for a subset of proteins uniquely found or differentially expressed (FDR<0.05, Fold change ≥ ±1.5) in the proteome of L-EV and S-EVs from a previously published study^8^. Characterization of the highly and differentially expressed proteins was performed by the Ingenuity Pathway Analysis (IPA, QIAGEN)^38^ and DAVID^39^ tools. Differentially expressed proteins among groups were selected based on an averaged relative expression difference >0.1 and a p-value<0.05 by one-sample t-test. Transcriptional information was recovered from the Prostate Cancer Transcriptome Atlas (PCTA; www.thepcta.org)^40^ and The Cancer Genome Atlas (TCGA, https://portal.gdc.cancer.gov)^41^ databases.

### Immunoblotting analysis

Immunoblotting analysis was performed as described^2^. Primary antibodies used were HSPA5 (#3177, 1:1,000 dilution), GAPDH (#3683, 1:10,000 dilution), H3 (#9717, 1:10,000 dilution) and pan-SRC (#2108, 1:10,000 dilution) from Cell Signaling. Antibodies for TSG101 (sc-7964, 1:1,000 dilution), CD9 (sc-13118, 1:1,000 dilution), STEAP1 (sc-271872, 1:1,000 dilution), and CAV1 (sc-894, 1:10,000 dilution) were obtained from Santa Cruz. Antibodies to KRT18 (ab93741, 1:10,000 dilution) and CD81 (ab79559, 1:10,000 dilution) were obtained from Abcam and ABCC4 (#GTX15602, 1:10,000 dilution) from GenTex. Antibodies to ACTB (A5441, 1:10,000 dilution) was obtained from Sigma and STEAP2 (#PA5-25495, 1:1,000 dilution) from Thermo Scientific. Densitometric quantification of films was performed with the ImageJ software (v1.52a).

### Tunable Resistive Pulse Sensing measurements (TRPS)

Concentration and particle size distribution of EVs were carried out in a qNano device (iZON Science, New Zealand). Freshly isolated EVs were diluted 1:40 in 0.2-µm filtered PBS and analyzed either with a NP2,000-nm nanopore for L-EVs or in a NP200-nm nanopore for S-EVs. Membranes were stretched at 47 mm and voltage set either at 0.04 V for L-EVs or 0.5 V for S-EVs in order to achieve a stable current baseline of about 120 nA. Particle size and concentrations were calibrated using Izon calibration particles (1:100 diluted TPK200 for S-EVs, and 1:1000 diluted CPC2000 for L-EVs) and a minimum of 500 events were registered for each sample with a positive pressure of 5 mbar. Particle quantitation performed in EVs obtained from ∼3.0×10^8^ cells and resuspended in 200 µL of filtered PBS.

### Flow cytometry (FC) analysis of prostate cancer cells and L-EVs

L-EVs were isolated as described above. Cells were fixed in 75% EtOH for 30 min at 4°C. L-EVs were fixed in 4% PFA for 10 min at room temperature. Cells and L-EVs were incubated for 30 min or 1 h, respectively, with one of the following primary antibodies: STEAP1 (H00026872-D01P, Abnova) at 1:50 dilution, STEAP2 (PA5-25495, Invitrogen) at 1:10 dilution, or ABCC4 (GTX15602, GeneTex) at 1:20 dilution. Excess antibody was washed off with PBS followed by incubation for 30 min with phycoerythrin-conjugated (111-116-144, Jackson ImmunoResearch; for STEAP1 and STEAP2) or FITC-conjugated (31629, ThermoFisher Scientific; for ABCC4) secondary antibody at 1:250 dilution. Both cell and L-EV samples were analyzed using the LSR-II flow cytometer (Becton Dickinson) with settings optimized for the detection of cells or particles larger than 1 µm, respectively. Data were analyzed using FlowJo software (Treestar).

### Statistical analysis

Plots represent the mean and standard deviation of at least three independent replicates. Experimental groups were compared using Student’s t test (unpaired, two-tails) and statistical significance established for a p-value<0.05.

## Results

### Low-background acyl-biotinyl exchange (LB-ABE) enables highly selective isolation of palmitoyl-proteins from whole cell lysates and membrane preparations

Because palmitoyl-proteins are anchored to cellular membranes^1^ and EVs are enriched in membrane components, we hypothesized that palmitoyl-proteins would be enriched in EVs. Using an *in silico* approach, we intersected the compendium of palmitoylated proteins SwissPalm^36^, with the ExoCarta database that contain proteins identified in EVs^37^. As expected, we found a 3-fold higher percentage of putative palmitoylated proteins in EVs versus cells (13.2% versus 4.3%) **(Suppl. Figure 1A)**. Further *in silico* analysis of the whole proteomes of prostate cancer cell-derived L- and S-EVs^8^, obtained by gradient centrifugation, showed that ∼20% of the proteins identified in both EV fractions are putative palmitoylated proteins **(Suppl. Figure 1B)**. Additionally, when we examined the proteins that are differentially expressed between the two EV fractions, this percentage increased to 43% in L-EVs and 32% in S-EVs **(Suppl. Figure 1B)**. These results suggest that palmitoylated proteins are enriched in EVs in comparison to cells and might be differentially distributed in different EV populations.

Quantitative analysis of protein palmitoylation has been impaired by the lack of methodological approaches allowing efficient separation of palmitoylated proteins from non-palmitoylated proteins. We recently developed a substantially improved acyl-biotinyl exchange (ABE) method, termed low-background ABE (LB-ABE)^2^, which consists of the blockage of non-palmitoylated cysteine residues by N-ethylmaleimide (NEM) and further by 2,2’-dithiodipyridine (DTDP), followed by converting palmitoylated cysteine residues into biotinylated cysteines and specific purification of palmitoyl-proteins by streptavidin **(Figure 1A).** This approach largely eliminates the co-isolation of non-palmitoylated proteins, thus enabling a comprehensive and specific palmitoyl-proteomic analysis^2^.

**Figure 1.**
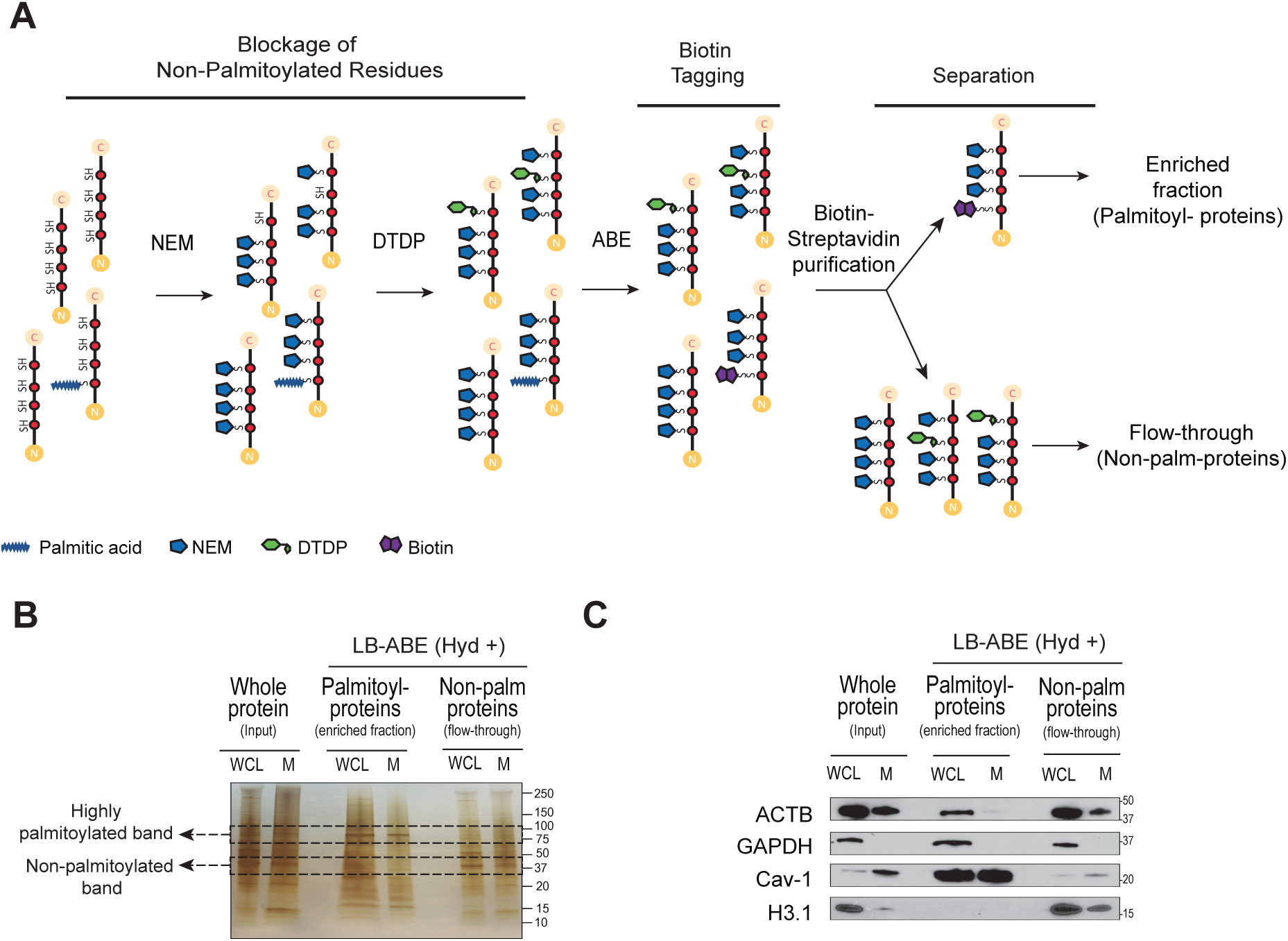
Selective enrichment of putative palmitoylated proteins in the proteome of EVs. **A)** Schematic representation of the low-background acyl-biotinyl exchange (LB-ABE) method employed for selective enrichment of palmitoylated proteins. Free cysteines are sequentially blocked by NEM and DTDP incubations. Acyl-biotinyl exchange (ABE) allows specific labeling of palmitoyl-proteins and purification of the whole palmitoyl-proteome by biotin-streptavidin interaction. Non-palmitoylated proteins can be recovered in the flow-through upon specific capture of palmitoyl-proteins with streptavidin-functionalized beads. **B)** Silver-stained PAGE gel of PC3 WCL and M protein lysates with versus without LB-ABE enrichment of palmitoyl proteins. **C)** Immunoblotting of select proteins enriched or excluded from the whole cell and membrane palmitoyl-proteome of PC3 upon LB-ABE.

We tested the LB-ABE method in membrane preparations (M) from PC3 prostate cancer cells before applying it to EVs because M preparations are easier to generate and give a higher protein yield than EVs. Silver staining of M processed through LB-ABE enrichment showed distinct protein patterns in the fractions enriched with or depleted of palmitoyl-proteins **(Figure 1B)**. This enrichment was further validated by immunoblotting for proteins known to exhibit variable palmitoylation status. Caveolin-1 (Cav-1), a membrane-associated protein, was highly enriched in the palmitoyl-protein fraction captured by LB-ABE **(Figure 1C)**, and almost completely absent from the palmitoylprotein-depleted fraction obtained from both M and WCL. This remarkable enrichment suggests that Cav-1 is predominantly present as a palmitoylated protein, in agreement with our recent report showing that Cav-1 is nearly 100% palmitoylated^2^. In contrast, nuclear histone H3 (H3.1), which is predicted to be a non-palmitoylated protein by the SwissPalm database, was completely excluded from the palmitoyl-protein fractions **(Figure 1C)**. Interestingly, GAPDH, which is a cytosolic protein and was not detected in M, was present in similar proportions in its palmitoylated and non-palmityolated form in WCL fractions, whereas another cytoplasmic protein, β-actin (ACTB), was mostly non-palmitoylated, even when detected in M, suggesting that palmitoylation is not essential for its membrane association. The succesfull enrichment of palmitoylated proteins by LB-ABE in WCL and M encouraged us to apply this approach to studying protein palmitoylation in EVs at a large scale.

### Comprehensive palmitoyl-proteomic analysis identifies differentially abundant palmitoyl-proteins in WCL, M, and EVs

In order to identify EV-specific palmitoyl-protein signatures, we employed liquid chromatography-tandem mass spectrometry (LC-MS/MS) to analyze palmitoylated proteins isolated by LB-ABE in L-EVs and S-EVs, and performed a comparative analysis with the palmitoyl-proteins identified in WCL and M from PC3 cells. L-EVs and S-EVs were isolated from cell culture media by differential ultracentrifugation followed by density gradient purification **(Suppl. Figure 1C)**, in line with the most recent MISEV2018 guidelines^11^ and previous studies that separated large oncosomes from exosomes^7,8,25,42^. Over 99% of the cells from which EVs were isolated were viable **(Suppl. Figure 1D)** limiting the possibility of contamination by apoptotic bodies.

Particle concentration and size distribution of L-EV and S-EV preparations were analyzed by TRPS, using a qNano particle analyzer (Izon Sciences) **(Figure 2A)**. L-EV samples contained 1.76 × 10^9^ particles/mL of 1.5-5 µm diameter, with a modal size of 1.77 µm. S-EV samples contained 2.43 x 10^11^ particles/mL of 90-600 nm diameter, with a modal size of 131 nm, in line with previous reports on L- and S-EVs of the type of large oncosomes and exosomes^7^. To further characterize L-EVs and S-EVs, we evaluated total protein yield and found that L-EVs contained significantly more protein than S-EVs normalized to the number of producing cells **(Figure 2B)**, in agreement with our previous observations^8^. Finally, immunoblotting of proteins enriched in L- and S-EVs confirmed the nature of the EV preparations (Figure 2C). L-EVs showed the enrichment of HSPA5 and KRT18 at 1.10-1.15 g/mL, which are enriched in large oncosomes^7,8,25,43^, while S-EVs showed the enrichment of CD81 and TSG101 at 1.10 g/mL, that are typically enriched in exosomes **(Figure 2C)**, in line with current literature^6–8,11,25,43^.

**Figure 2.**
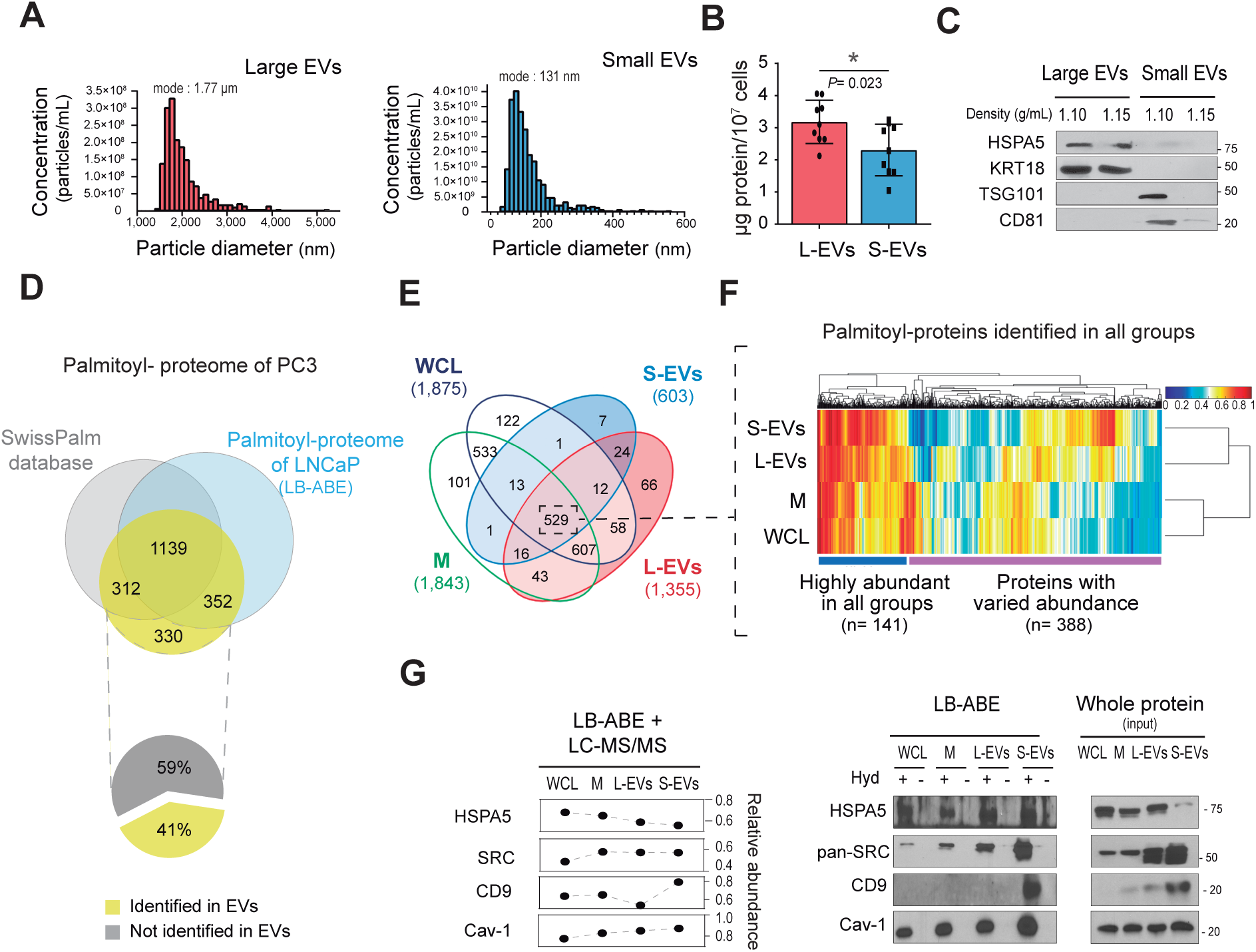
Large-scale MS analysis identifies the palmitoyl-protein signature of prostate cancer cells and EVs. **A)** Quantification and particle size distribution of PC3 EVs by TRPS. NP2000 (resolution window 0.9-5.7 µm) and NP250 (resolution window 110-630 nm) nanopores were used for the quantification of L- and S-EVs, respectively. Histogram plots depicted with a bin width of 100 and 10 nm, respectively. **B)** Yield of purified PC3 EV-protein after differential ultracentrifugation and density-gradient purification of conditioned media. **C)** Immunoblotting of select proteins enriched either in L-EVs (HSPA5 and KRT18) or S-EVs (TSG101 and CD81). **D)** Venn diagram showing the number and overlap of palmitoylated proteins identified in PC3 cells by LB-ABE (n=2,133) in comparison to LNCaP cells ^2^ and the number of known human palmitoyl-proteins compiled in the SwissPalm database^36^. **E)** Venn diagram showing the number of palmitoylated proteins identified in WCL, M, L-EVs and S-EVs and their level of overlap (n=2,339). **F)** Unsupervised heat map and dendrogram of the normalized relative abundance of the common palmitoyl-proteins identified (n=529). Clustering analysis identifies a group of palmitoylated proteins highly abundant across all subcellular compartments (n=141) and a group of proteins with varied abundance (n=388). **G)** Left, dot plot shows the relative abundance of select palmitoyl-proteins detected in WCL, M, L-EVs and S-EVs by MS analysis and max(1)-min(0) normalization. Middle, immunoblotting against the indicated palmitoyl-proteins from WCL, M, L-EVs and S-EVs in presence of hydroxylamine (Hyd+) or not (Hyd-) confirms the specific enrichment of palmitoyl-proteins by LB-ABE. Right, immunoblotting of 2 µg of WCL, M, L-EVs and S-EVs taken prior the enrichment of palmitoylated proteins in order to confirm the expression and distribution of the indicated proteins.

Next we compared the palmitoyl-proteins identified in WCL, M, L-EVs, and S-EVs. A total of 2,408 palmitoyl-proteins were identified with a false discovery rate (FDR) of <0.01. We then selected 2,133 high confidence proteins detected in two or more replicates and no less than 5% of absolute protein abundance **(Suppl. Table 1)**. Of these, 1,803 proteins are classified as palmitoyl-proteins in the SwissPalm (v2) database^36^ and/or were identified in our recent deep palmitoyl-proteomic profiling of prostate cancer LNCaP cells^2^ **(Figure 2D)**. Of the remaining 330 proteins, which have not yet been reported as palmitoyl-proteins, 41% were identified in EVs. This result confirms successful enrichment of palmitoyl-proteins with the LB-ABE approach, which outperforms conventional ABE methods, thus allowing the identification of novel palmitoyl-proteins, including in EV fractions.

Specifically, a total of 1,875, 1,843, 1,355, and 603 palmitoyl-proteins were identified in WCL, M, L-EVs, and S-EVs, respectively **(Figure 2E and Suppl. Table 1)**. The reason why a much smaller number of proteins were identified in S-EVs is that lowly abundant species not detected in at least two replicates were removed, and only high-confidence proteins with high correlation among replicates were retained for further analysis **(Suppl. Figure 1E).** Clustering of the palmitoyl-proteins detected in all 4 fractions (WCL, M, L-EVs, and S-EVs) (n=529) **(Figure 2E)** identified 141 proteins that were consistently highly abundant (see Figure legend) across all fractions and 388 proteins with varied abundance **(Figure 2F)**. Among the proteins with varied abundance, 31 were more abundant in L-EVs and 98 in S-EVs **(Suppl. Table 1)**, suggesting that S-EVs harbor a more discrete palmitoyl-protein signature than L-EVs. Functional enrichment analysis of the 141 highly abundant proteins identified in all four fractions including EVs showed significant association with major biological processes relevant to cancer (e.g., adhesion and cellular movement) **(Suppl. Figure 2A)**. Importantly, vesicle-mediated transport was also identified as one of the most prominent functions of the identified palmitoylated proteins **(Suppl. Figure 2A)**. These data suggest functional significance of palmitoylation in the biology of cancer and EVs. Importantly, they corroborate the utility of this post-translational modification analysis as a means for discovering novel EV biomarkers.

Western blotting confirmed expression of selected palmitoyl-proteins in the fractions enriched for palmitoylated proteins (Hyd+) but not in the control groups (Hyd-) confirming specific recovery following LB-ABE **(Figure 2G)**. Src-family tyrosine kinases and Cav-1 are known to be highly palmitoylated in prostate cancer^44,45^, and they were identified with high confidence in all fractions **(Figure 2G)**. HSPA5 is enriched in L-EVs as total protein^7,8,25,43^ but we found it to be equally represented in L- and S-EVs as a palmitoylated protein. CD9 is enriched in S-EVs both as a total and as palmitoyl-protein. We also noticed that the palmitoylated form of CD81 was significantly enriched in L-EVs **(Suppl. Table 1** and data not shown), despite its “canonical” description as an S-EV marker^6–8,11,25,43^, based on total protein analysis. This result suggests that the palmitoyl status of certain proteins can alter their enrichment in different EV fractions.

### L- and S-EVs exhibit distinct profiles that distinguish them from their parental cells

We and others have previously demonstrated that EVs contain not only proteins that are highly expressed in the originating cells, but they are also enriched in proteins of low abundance^21^. Moreover, discrete EV populations contain a set of distinct proteins, suggesting cargo selection^6,8^. In order to determine whether this cargo selection is evident in the palmitoyl-proteome, we compared the content of L- and S-EVs to the originating cells. The palmitoyl-proteome of WCL and M fractions were strongly correlated (r=0.782) suggesting that our method efficiently identifies palmitoyl-proteins even in WCL. Because palmitoylation is a membrane-anchoring modification and EVs are membrane encapsulated particles, we compared the palmitoyl proteins identified in EVs with those identified in M fractions. The majority (roughly 90%) of the palmitoyl-proteins identified in both L- and S-EVs were also identified in M **(Suppl. Figure 2B)**, demonstrating the overall similar composition of the cellular and EV membranes. However, when we looked at relative abundance of these common proteins, and compared their expression in M versus both EV types, we found a weak correlation (r=0.444 in L-EVs; r=0.317 in S-EVs) **(Figure 3A)**, suggesting selective enrichment of palmitoylated proteins in EVs. Functional enrichment analysis identified a series of cell functions overrepresented in EVs in comparison to M. More specifically, the palmitoyl-proteins significantly enriched in both L- and S-EVs were involved in pyridine nucleotide metabolism and cell adhesion **(Figure 3B and Suppl. Table 2)**, including proteins known to be involved in modulating cell polarity, adhesion and migration^46^. These comprised membrane-anchored integrins (ITGA2, ITGA6, ATGB4), claudins (CLDN1, CLDN1), and cytoskeletal proteins such as the Rho-associated moesin (MSN), vinculin (VCL), ezrin (EZR), ROCK2 and the Ras GTPase-activating-like protein IQGAP1 **(Suppl. Table 2)**. Cell functions overrepresented in proteins more abundant in L-EVs than in M were associated with positive regulation of organelle and cell component organization, as well as actin cytoskeleton organization **(Figure 3B, Suppl. Table 2)**. In contrast, proteins associated with cell and chemical homeostasis were over-represented in S-EVs **(Figure 3B, Suppl. Table 2)**.

**Figure 3.**
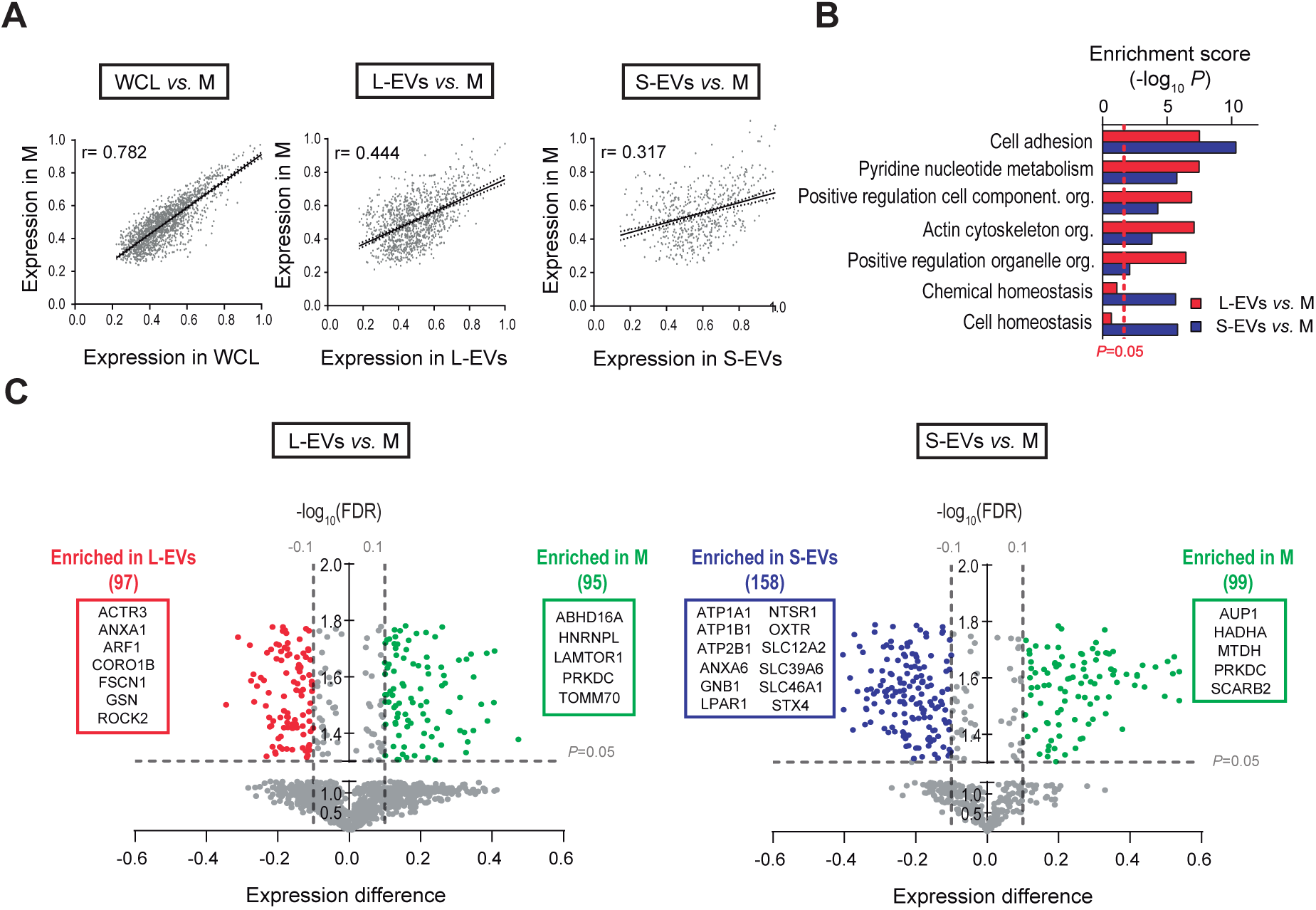
L- and S-EVs exhibit distinct palmitoyl-protein profiles that distinguish them from their parental cells. **A)** Regression analysis of the relative abundance of the palmitoyl-proteins identified in WCL, M, L-EVs and S-EVs. Spearman’s coefficient (r) demonstrates a low correlation between EVs and M when compared to WCL and M. **B)** The biological functions overrepresented in L- and S-EVs in comparison to M identified by functional enrichment analysis of the palmityol-proteome differentially expressed in EVs using DAVID software^39^. **C)** Volcano plots showing differential protein expression between L- and S-EVs compared to M. X and Y axes represent the normalized expression difference and -log_10_(FDR), respectively. Red and blue dots correspond to palmitoyl-proteins significantly enriched either in L- or S-EVs, respectively, compared to those enriched in the M (green dots). The blue and red boxes highlight functionally relevant proteins to the biological processes differentially represented in L-EVs and S-EVs shown in panel B. The green boxes highlight the top 5 proteins enriched in M.

Detailed analysis of the palmitoylated protein species involved in these biological processes revealed that, while the high-abundance proteins in L-EVs were mostly represented by cytoplasmic proteins involved in cell growth (GSN, FSCN1 and ATCR3) and signal transduction (ANXA1, ARF1, ROCK2 and CORO1B); the proteins enriched in S-EVs were mainly represented by two major groups of plasma membrane proteins **(Figure 3C)**. The first group contained two families of transmembrane palmitoyl-proteins with transporter activity: the P-type ATPase superfamily (ATP1A1, ATP1B1, ATP2B1) and the cation transporters (SLC12A2, SLC39A6, and SLC46A1). The second group contained palmitoyl-proteins associated with signal transduction and cell communication (ANXA6, LPAR1, GNB1, LYN, NTSR1, OXTR and STX4) **(Figure 3C)**.

### Palmitoyl-proteomic profiles of L-versus S-EVs reflect EV population-specific biological processes and subcellular origin

To further explore the difference between the palmitoyl-proteomes of L-EVs and S-EVs, we performed comparative analysis of the proteins uniquely or differentially expressed in each EV population **(Suppl. Table 1)**. Ninety-seven palmitoyl-proteins were detected only in EVs, with 66 unique to L-EVs, 7 to S-EVs, and 24 shared EV in both EV types) **(Figure 2E)**. We also identified 581 proteins (24+12+529+16) that were detected in both S-EV and L-EVs but not unique to them (**Figure 2E**). Among them, 146 proteins were significantly enriched in L-EVs and 151 proteins were significantly enriched in S-EVs **(Figure 4A)**. Functional analysis showed that palmitoyl-proteins enriched in L-EVs were associated with protein localization, regulation of protein stability, regulation of cell component organization and cellular localization **(Figure 4B)**. This association was mainly represented by the Vacuolar Protein-sorting Associated Protein 35 (VPS35), which is involved in sorting of proteins via the cargo-selective complex (CSC)^47^, and by several members of the chaperonin-containing T-complex (CCT2, CCT3, CCT4, CCT7), which have been described to participate in the folding of nascent proteins^48^, as well as in vesicular transport within the cell^49^ **(Figure 4A)**. Functional analysis of the palmitoyl-proteins enriched in S-EVs showed enrichment for proteins associated with cell communication, signaling processes, proteolysis and regulation of phosphate metabolism **(Figure 4B)**, and included metalloproteinase ADAM17, the death receptor FAS and the ubiquitin UBA52 **(Figure 4A)**. The palmitoylated form of CD9, a tetraspanin highly enriched in S-EVs, was identified as significantly enriched in S-EVs in comparison to L-EVs **(Figure 4A)** – in agreement with the distribution observed for the non-palmitoylated form.

**Figure 4.**
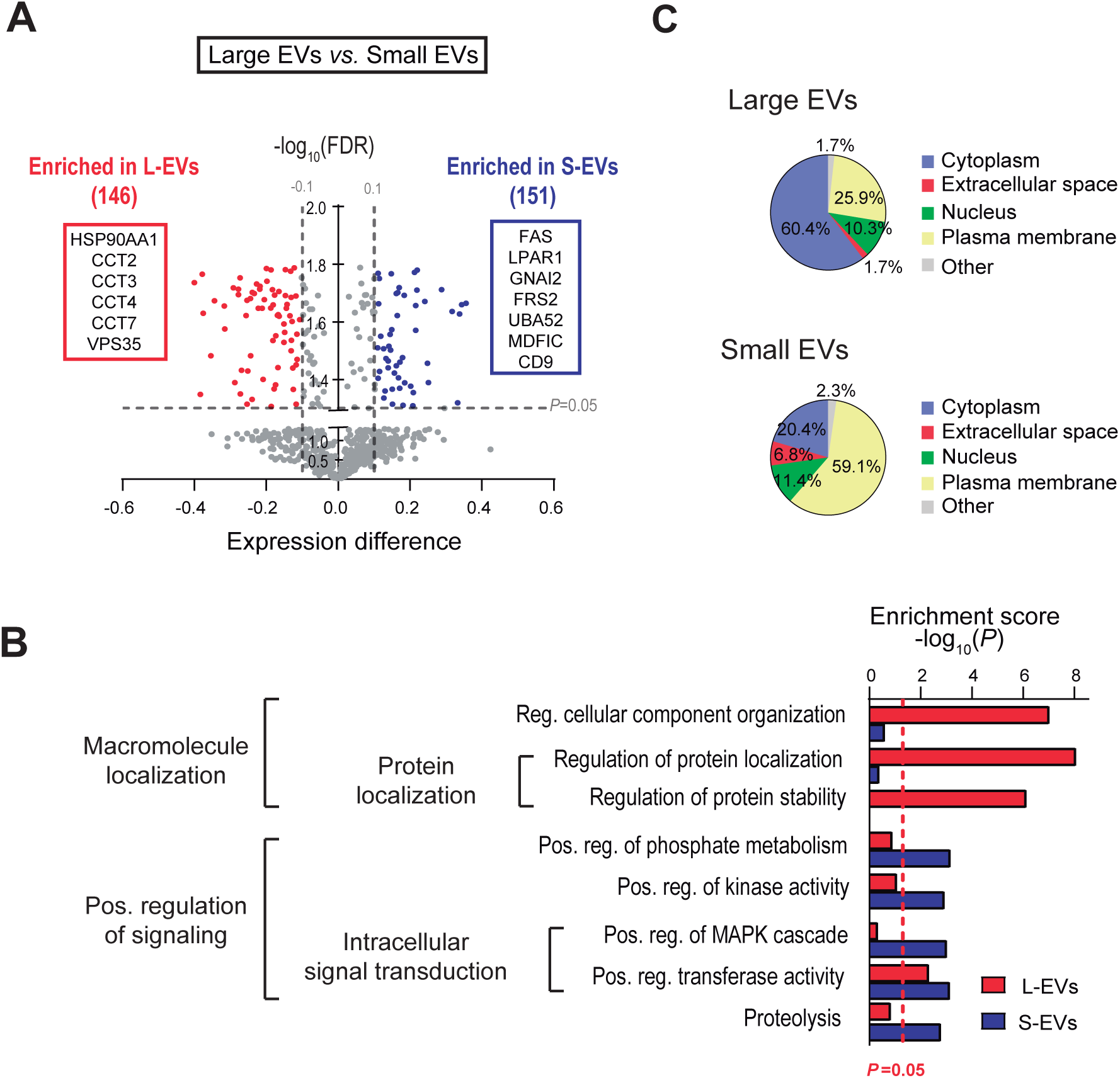
Palmitoyl-protein profiles of L- and S-EVs are associated with EV population-specific biological processes and subcellular origin. **A)** Volcano plot showing the differential protein expression between L-EVs and S-EVs. Red and blue dots correspond to proteins significantly enriched in L- and S-EVs, respectively. X and Y axes represent the normalized expression difference and -log_10_(FDR), respectively. The red and blue boxes highlight functionally relevant palmityol-proteins to the biological processes differentially represented in L- and S-EVs shown in panel B. **B)** The biological functions overrepresented in L- and S-EVs identified by functional enrichment analysis of the palmitoyl-proteome differentially expressed in EVs using DAVID software. **C)** Pie charts indicating the subcellular localization of the palmitoylated proteins, as defined by the Ingenuity Knowledge database, differentially expressed in L- (n=58) and S-EVs (n=44).

Next we examined subcellular distribution of the palmitoyl-proteins differentially expressed in L- and S-EVs. Palmitoyl-proteins enriched in L-EVs were primarily associated with cytoplasm (60.4%), while those enriched in S-EVs were mainly associated with the plasma membrane (59.1%) **(Figure 4C)**. Interestingly, both EV populations contained a smaller portion of nuclear proteins, which were similarly represented in both EV types (10.3% in L-EVs, 11.4% in S-EVs). Altogether, these data suggest a different subcellular derivation and functional/biological profiles of the palmitoylated proteins in L- and S-EVs.

### Prostate cancer-derived EVs contain cancer-specific palmitoylated proteins

In order to determine if the protein palmitoylation profile in EVs reflected cancer-associated functions, we performed functional analysis of the palmitoylated proteins abundant in EVs **(Suppl. Table 1)**. We found an association with canonical cancer functions such as cell-to-cell signaling, adhesion, cell movement and EMT **(Figure 5A)**.

**Figure 5.**
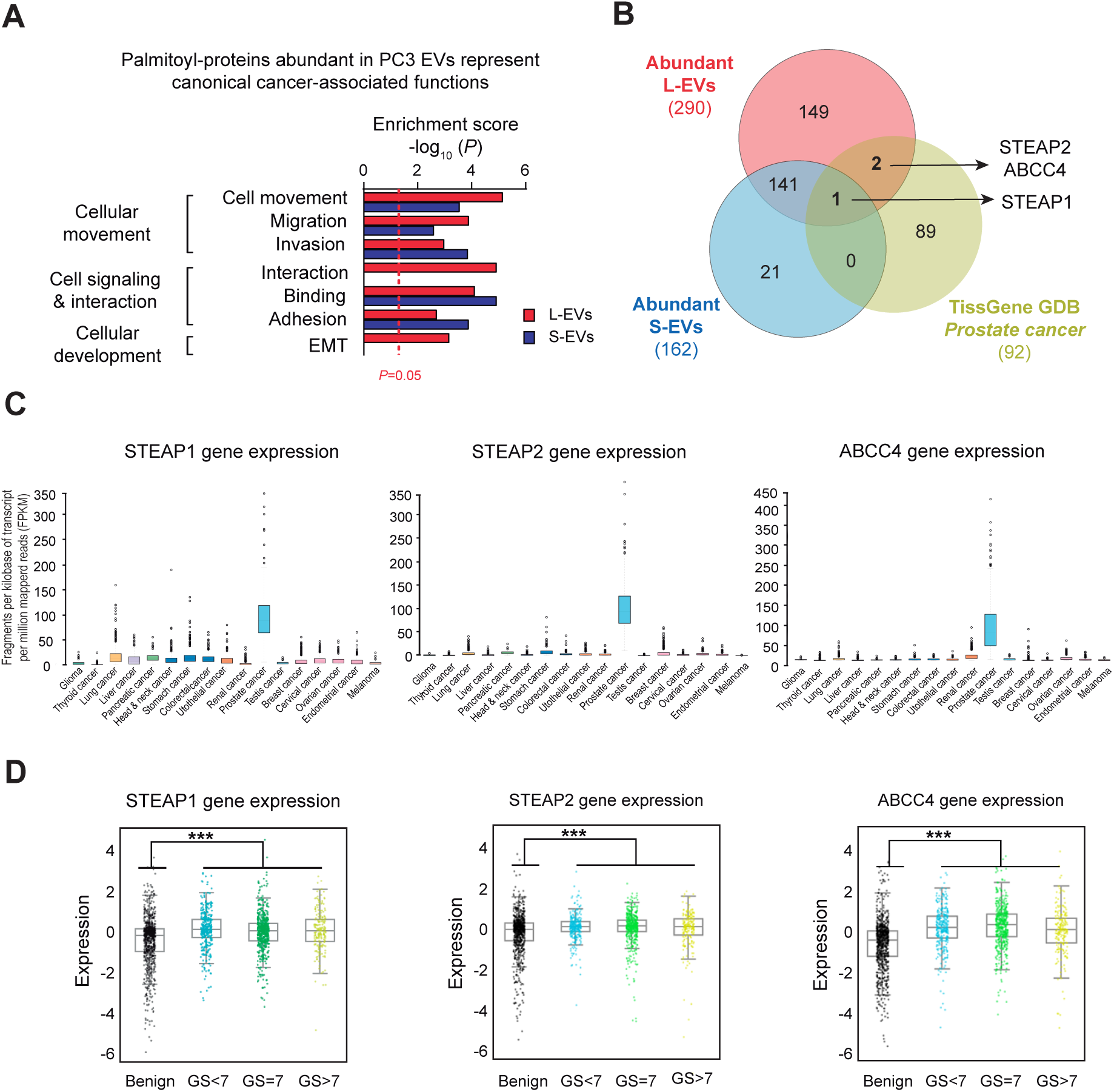
Prostate cancer-derived EVs are enriched in prostate cancer-enriched palmitoyl-proteins. **A)** Cancer-associated functions of the palmitoyl-proteins abundant in L- and S-EVs identified using IPA. **B)** Venn diagram showing the number of palmitoyl-proteins abundant (determined by the rank product algorithm) in L- and S-EVs which are specifically enriched in prostate cancer according to the TissGene GDB^50^. **C)** The gene expression of STEAP1, STEAP2 and ABCC4 determined as fragments per kilobase of transcript per million mapped reads (FPKM) for select carcinomas according to the cancer genome atlas (TCGA) database^41^. **D)** The change in gene expression of STEAP1, STEAP2 and ABCC4 with prostate cancer progression according to the prostate cancer transcriptome atlas (PCTA) database^40^.

Using a panel of tissue-specific genes that combines the information from different databases^50^, we identified 92 prostate cancer-specific/enriched genes. We identified the multidrug-resistance-associated protein 4 (ABCC4), and the six-transmembrane epithelial antigen of prostate 1 (STEAP1) and 2 (STEAP2) **(Figure 5B)** as abundant palmitoyl-proteins in EVs. Importantly, these 3 genes are codified by RNA that is highly expressed in prostate cancer tissue in comparison to other types of cancer **(Figure 5C)** and benign prostate lesions **(Figure 5D)**. Interestingly, even though these proteins were abundant in both EV types, palmitoylated STEAP1 was more abundant in S-EVs, while STEAP2 and ABCC4 were more abundant in L-EVs **(Suppl. Figure 2C)**, confirming that EV heterogeneity is not limited to size but also to cargo.

Immunoblotting confirmed strong expression of STEAP1 in both populations of EVs from highly metastatic PC3 cells in comparison with the cells themselves **(Figure 6A)**. This was not the case for purified EVs from DU145^DIAPH3-KD^ cells **(Suppl. Figure 2D-E)**, which is also highly metastatic, suggesting that STEAP1 expression in EVs is not correlated with disease aggressiveness **(Figure 6A)**. Conversely, STEAP2 was equally abundant in cells and both EVs from PC3 and DU145^DIAPH3-KD^ cells **(Figure 6B)**. Finally, ABCC4 was highly enriched in EVs from PC3 and DU145^DIAPH3-KD^ cells in contrast to the low expression detected in the originating cells **(Figure 6C)**. We also analyzed these proteins at the single EV level using flow cytometry, and found that STEAP1 was highly expressed in both PC3 and DU145^DIAPH3-KD^ L-EVs while found to be virtually undetectable in PC3 cells **(Figure 6D)**. STEAP2 was highly abundant in both L-EVs and cells **(Figure 6E)**. ABCC4 was relatively enriched in L-EVs versus cells **(Figure 6F)**. Collectively these results indicate that prostate cancer-specific proteins enriched in EVs are not necessarily identifiable in cancer tissue and support the use of EV cargo as a source of clinically relevant biomarkers.

**Figure 6.**
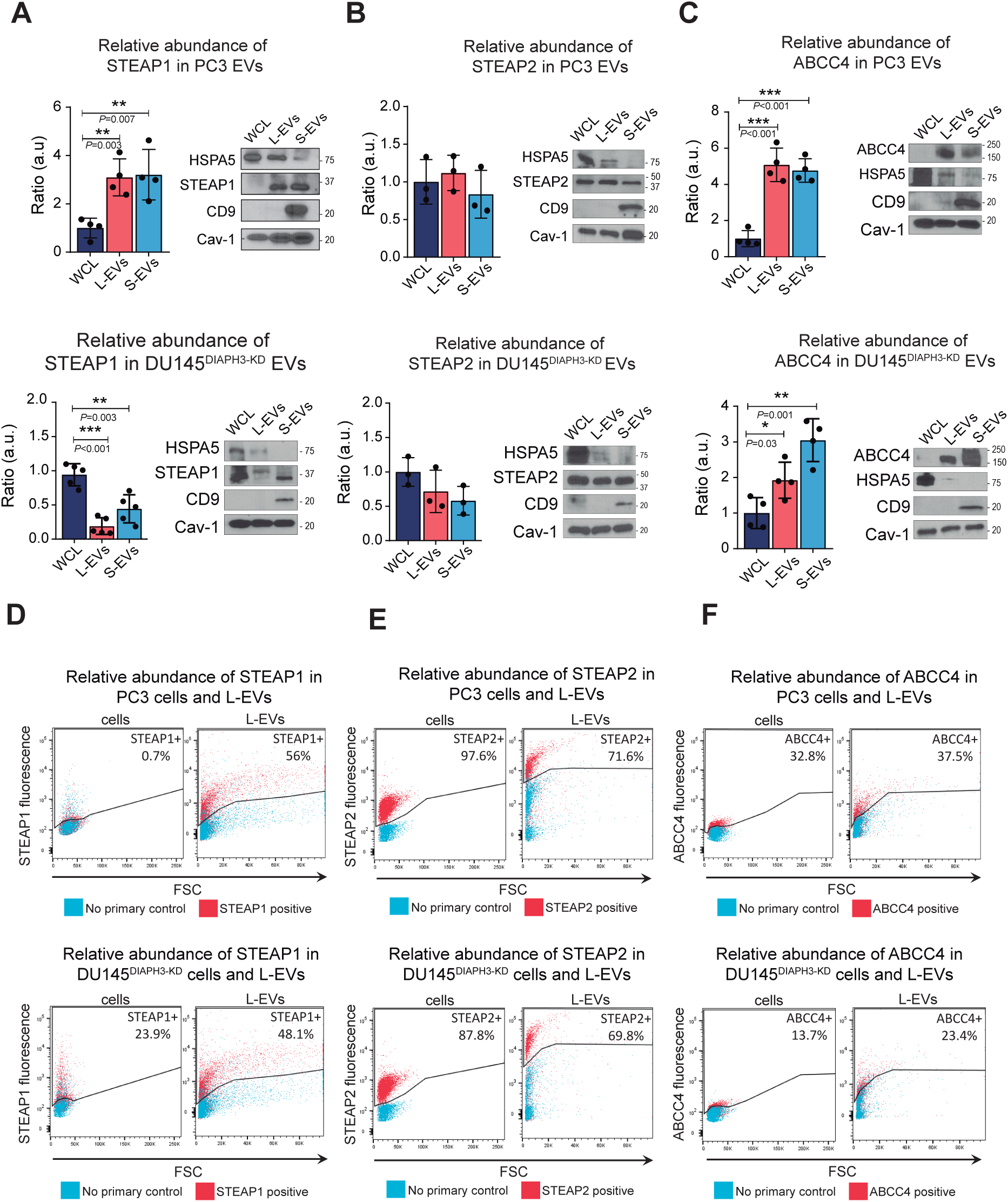
Prostate cancer-specific proteins are enriched in EVs. **A-C)** Representative immunoblot of STEAP1 (A), STEAP2 (B) and ABCC4 (C) in EVs and WCL from PC3 and DU145^DIAPH3-KD^ cells along with the control proteins HSPA5 (L-EV enriched protein), CD9 (S-EV enriched protein), and Cav-1 (loading control). Bar plots represent the densitometric quantification across replicate blots (n=3-5). **D-F)** FACS analysis of STEAP1 (D), STEAP2 (E) and ABCC4 (F) expressionin cells and L-EVs from PC3 and DU145^DIAPH3-KD^ cells.

Finally, we investigated if palmitoylation had any influence on the localization of select proteins in EVs. Treatment with the general inhibitor of protein palmitoylation 2-bromopalmitate (2BP) induced a general decrease of STEAP1 **(Figure 7A)**, STEAP2 **(Figure 7B)** and ABCC4 **(Figure 7C)** in EVs, suggesting a role for protein palmitoylation in the trafficking and sorting of these proteins to EVs. The 2-BP was used at a dose of 10 µM, which was the highest concentration at which cells remained completely viable **(Suppl. Figure 2F)**, and that did not alter the recovery of EV-protein **(Figure 7D)**. As expected, Cav-1 levels in EVs did not change as a consequence of 2-BP treatment **(Suppl. Figure 2G)**. This is a novel observation but in line with previous studies that report that palmitoylation of Cav-1 is not necessary for its membrane localization^51^.

**Figure 7.**
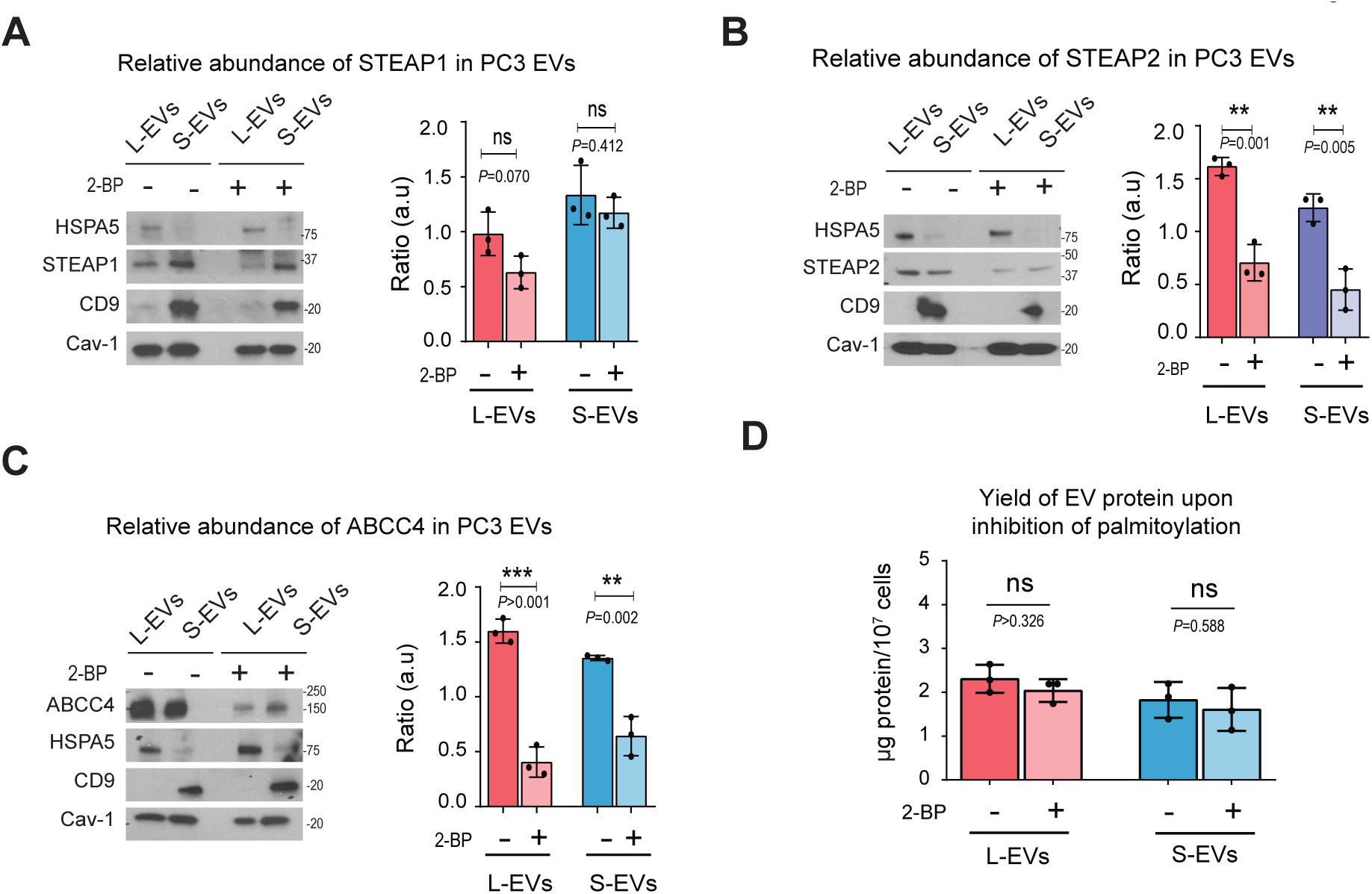
Inhibition of palmitoylation reduces the abundace of prostate cancer-specific palmitoyl-proteins in EVs. **A-C)** Representative immunoblot of the abundance of STEAP1 (A), STEAP2 (B) and ABCC4 (C) along with the control proteins HSPA5 (L-EV enriched protein), CD9 (S-EV enriched protein), and Cav-1 (loading control) in PC3 L- and S-EVs after inhibition of palmitoylation with 10 µM 2-bromopalmitate for 24h. Bar plots represent the densitometric quantification across replicate blots (n=3). **D)** Yield of purified EV-protein from PC3 cells treated with or without 10 µM 2-bromopalmitate for 24h.

## Discussion

Here we describe the first study to profile the palmitoyl proteome of EVs, prompted by the observation that EVs contain at least twice the number of palmitoylated proteins in comparison to the cellular proteome. While some of the palmitoyl proteins recapitulated expression of total protein in specific compartments (i.e. Cav-1 and SRC, which were present in all compartments and CD9, which was specific to S-EVs, both as a total and palmitoylated protein), other proteins show differential enrichement of the palmitoyl form in specific EV populations in comparison with total protein. HSPA5, for example, which is known to be enriched in L-EVs as total protein^7,8,43^, was equally represented in L- and S-EVs as a palmitoylated protein. CD81, which is described as a canonical S-EV marker, as total protein^6–8,11,25,43^, was significantly enriched in L-EVs, as a palmitoylated protein. Palmitoylation might thus be involved in the protein cargo sorting to different populations of EVs, and this could have important functional implications that would be missed if only the total protein is investigated. Further studies on the palmitoyl-proteome of L- and S-EVs may identify mechanisms that could be exploited to target specific proteins to, or to deplete specific proteins from, L- or S-EVs.

The selective packaging of palmitoyl-proteins in EVs is also supported by the weak correlation between expression levels of palmitoyl-proteins of both L- and S-EVs and the total cell membrane. This is a finding in line with what was previously reported in studies comparing the proteomes of EVs vs. originating cells at the total protein level^21^. Our results are also in line with a recent study demonstrating a poor correlation between protein expression levels in L- and S-EVs incomparison to their cell of origin^21^. In that study, as in the current one, the differences between EVs and cell of origin are more pronounced in the S-EV than L-EV cargo, in line with a different origin of these vesicles from distinct intracellular compartments^12,13^. The fact that these differences are reflected at the palmitoyl-proteome level suggests that palmiltoylation might be a critical step in protein sorting to EVs and/or EV biogenesis. To investigate that question in the future, comparative studies of the palmitoyl and total proteome of EVs and cells of origin will be essential to identify abundant palmitoyl-proteins that are potentially involved in these functions. With improved mass spectrometry technologies, the number of proteins identified in EV preparations has significantly increased. However, only a few studies have separated EVs from other particles or protein complexes prior to mass spectrometry. Therefore, public proteomic data contain a number of proteins that are associated with EVs because they are pulled down all together by ultracentrifugation. In fact, it is now evident that characterizing the protein profile of EVs by mass spectrometry requires important steps, such as separating EVs of different size by ultracentrifugation, after eliminating cells and cell debris, by applying centrifugation gradients that separate EVs based on their density^6,8,11,25^. Our data demonstrates that by using this approach we can deplete non-EV associated contaminants such as lipoproteins or albumin^52^ that are highly abundant in proteomics from crude EV preparations.

Almost 20% of the proteins identified as putative palmitoylated proteins in our study, using our highly sensitive LB-ABE method, are novel palmitoylated proteins that are not present in either the SwissPalm database (release 2, Feb, 2018)^36^ or a recent report that provides the most comprehensive analysis of palmitoylated proteins^2^. Of these novel proteins, 41% are present in EVs. This increased sensitivity of the LB-ABE method may allow for the identification of novel disease biomarkers from biological fluids. Attempts to identify disease biomarkers from biological fluids by traditional proteomics have been hampered by the huge background signals from enormously abundant proteins such as albumin and IgGs, which severely masks MS detection of low abundance but physiologically important proteins. Because most background proteins are water-soluble and not palmitoylated, palmitoyl proteomics with LB-ABE may allow for drastic depletion of high-abundance biofluid proteins and thus better detection of circulating disease biomarkers.

Another novel observation in our study points to the association to different subcellular compartments for palmitoyl proteins enriched in L-versus S-EVs, with an enrichment for proteins associated with cytoplasm in L-EVs and for proteins associated with the plasma membrane in S-EVs. This likely reflects that L-EVs are often generated as a result of membrane blebs inflated by the cytoplasm. In contrast, the enrichment of proteins associated with the plasma membrane is likely due to the high membrane/cytoplasm ratio in S-EV. Interestingly, a similar percentage of non-histone nuclear proteins was identified in both EV types. This is puzzling considering the accepted notion, based on whole-proteome analyses, that nuclear proteins are barely present in EVs and in particular in S-EVs. The absence of these nuclear proteins in S-EV may be due to the use of bulk proteomics. It is possible that the abundance of the nuclear proteins is several orders of magnitude lower than that of most abundant EV proteins, so they could not be detected by whole EV-proteome analysis. However, by employing palmitoyl proteomics, many abundant EV proteins are depleted, so we are able to identify these low abundant nuclear proteins. These nuclear proteins include several proteins that have been proposed as drivers and liquid biopsy markers for cancer including CSE1L^53^ and HUWE1^54^. Furthers studies to understand how these proteins are incorporated into EVs may yield important insights into cancer progression. Interestingly, S-EVs contain the palmitoylated form of the palmitoyl-transferase ZDHHC5, which palmitoylates EZH2, one of the major transcription factors in prostate cancer progression^55^. This suggests that prostate cancer S-EVs may promote the transformation of other cells by inducing horizontal EV transfer of EZH2; however, further investigation is required. It is important to keep in mind that many nuclear proteins shuttle across different compartments, therefore the gene ontology enrichment analysis might not reflect the actual subcellular localization of these palmitoylated proteins. In line with the differences in subcellular compartments from which the EV cargo are derived from, there are also differences in the predicted functions of L- and S-EVs. L-EVs are enriched in proteins associated with protein localization and stability, whereas S-EV are enriched in proteins associated with metabolism and cellular homeostasis. Together this suggests that different populations of EVs have distinct origins and functions.

We found three proteins that are abundant in EVs (STEAP1, STEAP2, ABCC4) whose expression has been reported to be specific to prostate cancer. While STEAP1 was enriched in S-EVs, STEAP2 was found to be equally abundant in S- and L-EVs. The observation that STEAP2 participates in the intracellular vesicular transport machinery and associates with endocytic and exocytic pathways^56,57^ might explain its equal abundance in both L- and S-EVs. Both STEAP1 and ABCC4 were enriched in EVs in comparison to cells in aggressive prostate cancer cells when we examined the total protein. This is an important result, suggesting that proteins released by cancer cells are not necessarily the most abundant cell cargo. This confirms their importance and paves the way for translational studies aimed to validate this findings in patient biological fluids.

In summary, this is the first large-scale analysis of the palmitoyl-proteomes of S- and L-EVs. We showed 1) that the LB-ABE method enables the isolation of palmitoyl-proteins from whole cell lysates and membrane preparations, as well as from L- and S-EVs, 2) that L- and S-EVs exhibit EV population-specific profiles that reflect EV biological processes and subcellular origin and distinguish them from their parental cells; 3) that prostate cancer-derived EVs contain cancer-specific palmitoylated proteins. Altogether, our results suggest that palmitoylation may be involved in the sorting of proteins to different EV populations and may allow for better detection of disease biomarkers.

## Geolocation

Cedars-Sinai Medical Center, Los Angeles, California.

## Acknowledgements

The authors are grateful to the Cedars-Sinai flow cytometry core for experiments on extracellular vesicles, and to Drs. Leonora Balaj, Valentina Minciacchi, and Cristiana Spinelli for helpful discussions.

## Figure legends

**Supplemental Figure 1.**
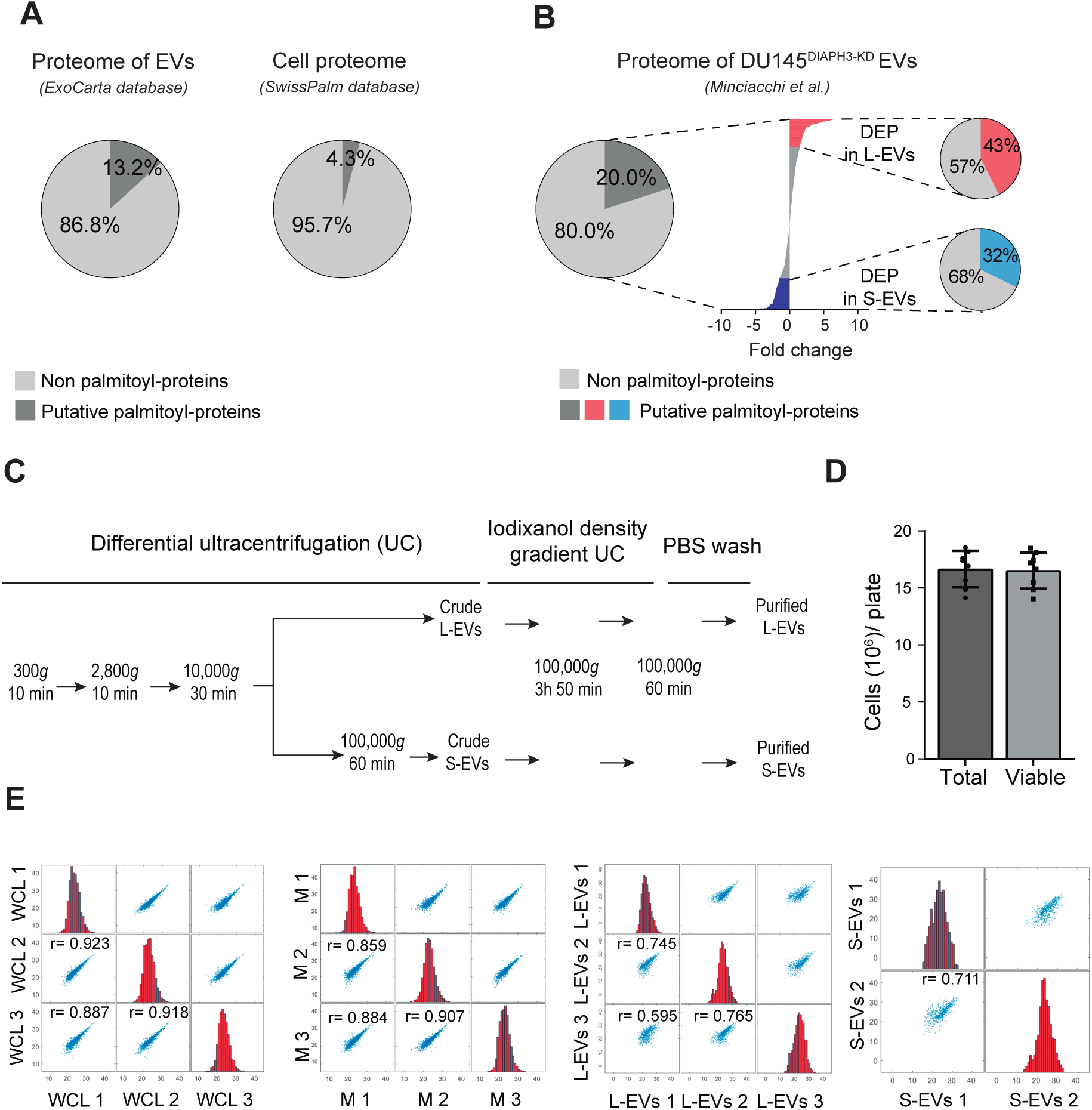
**A)** The left pie chart indicates the percentage of human proteins identified in EVs (n=5,403) according to the ExoCarta database, that are validated putative palmitoylated proteins, according to the SwissPalm database (n=972). The right pie chart indicates the percentage of validated palmitoyl-proteins in the human proteome (n=22,456), according to the SwissPalm database. **B)** (Left) The percentage of L- and S-EV proteins from our previously published study^8^ which have been reported to be palmitoylated in the SwissPalm database. (Right) The percentage of differentially expressed proteins (DEP)(fold change >±1.5 of the log_2_-transformed LFQ signal) in L- (n=90) and S-EVs (n=94) from the human prostate cancer cell line DU145^DIAPH3-KD^ which have been reported to be palmitoylated in the SwissPalm database. **C)** Schematic diagram for the isolation and purification of EVs. **D)** The average number of PC3 cells in a 150cm^2^ culture plate after 24 hours of FBS starvation. **E)** (Blue) Spearman’s correlation plot and coefficient (r) of the normalized LFQ signal among technical replicates in WCL, M, L-EVs and S-EVs (p-value<0.001). (Red) Histogram of the normalized LFQ signal for each replicate.

**Supplemental Figure 2.**
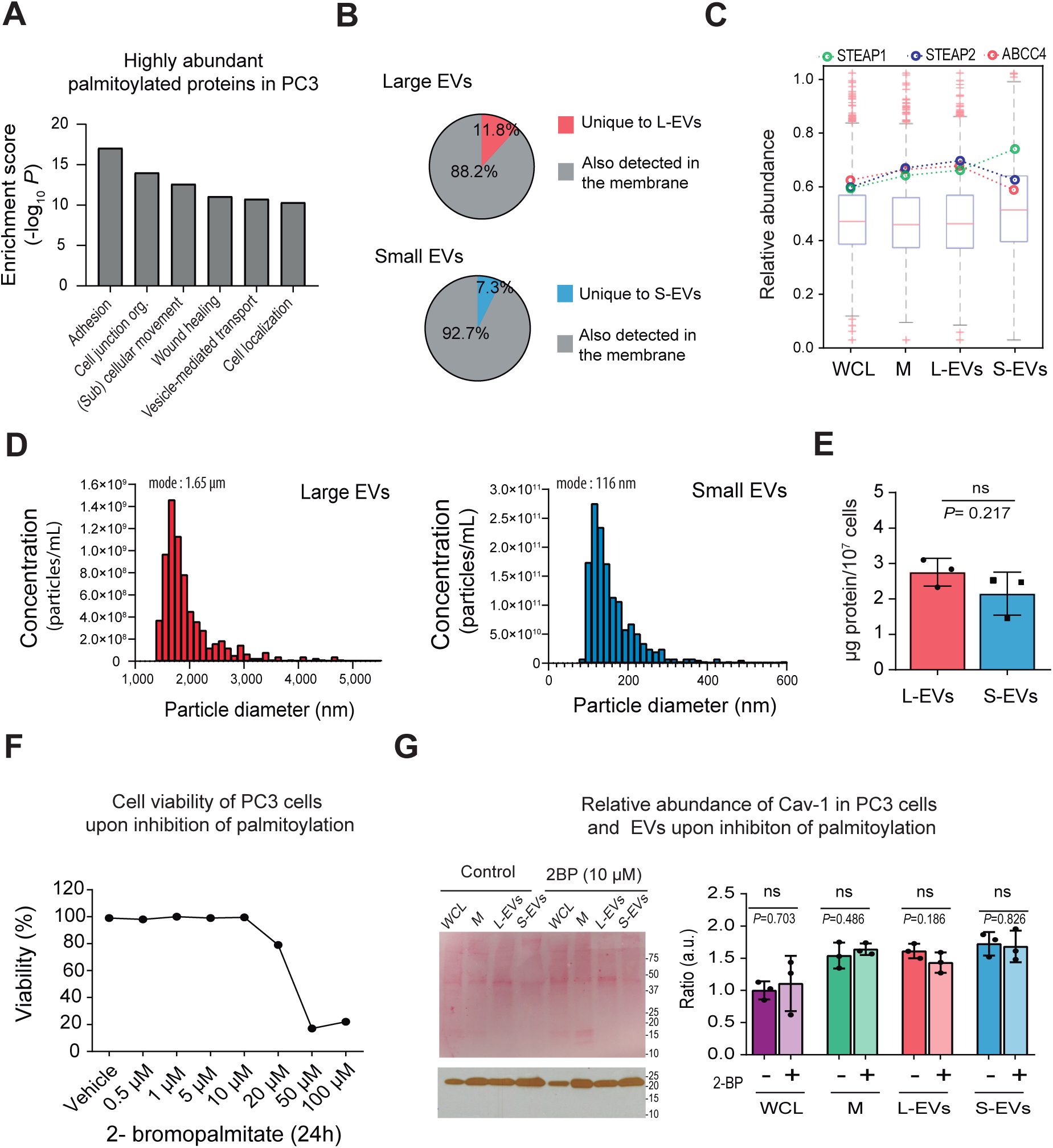
**A)** Functional enrichment analysis of the palmityolated proteins abundant (determined by the rank product algorithm) in PC3 cells and EVs (n=141) using DAVID software. **B)** The percentage of palmitoylated proteins identified in L- (n=1195) and S-EVs (n=559) that were identified in the palmitoyl-proteome of M in PC3 cells. **C)** The relative abundance of palmitoylated proteins detected across the indicated fractions by MS analysis. Dots indicate the relative abundance of the palmitoylated form of STEAP1 (green), STEAP2 (blue) and ABCC4 (red). **C)** Quantification and particle size distribution of DU145^DIAPH3-KD^ EVs by TRPS. NP2000 (resolution window 0.9-5.7 µm) and NP250 (resolution window 110-630 nm) nanopores were used for the quantification of L- and S-EVs, respectively. Histogram plots depicted with a bin width of 100 and 10 nm, respectively. **D)** Yield of purified DU145^DIAPH3-KD^ EV-protein after differential ultracentrifugation and density-gradient purification of conditioned media. **E)** Dot plot of the percentage of viability of PC3 cells after treatment with 2-bromopalmitate for 24 hours at the indicated doses. **F)** (Left) Ponceau staining and immunoblotting of Cav-1 in PC3 cells and EVs obtained upon inhibition of palmitoylation with 10 µM 2-bromopalmitate for 24h. L-EVs. (Right) Bar plots represent the densitometric quantification of Cav-1 across replicate blots (n=3).

